# Classifying drugs by their arrhythmogenic risk using machine learning

**DOI:** 10.1101/545863

**Authors:** Francisco Sahli Costabal, Kinya Seo, Euan Ashley, Ellen Kuhl

**Affiliations:** Department of Mechanical Engineering, Stanford University, Stanford, CA 94305, USA; Department of Medicine, Stanford University, Stanford, CA 94305, USA; Department of Pathology, Stanford University, Stanford, CA 94305, USA; Department of Bioengineering, Stanford University, Stanford, CA 94305, USA

## Abstract

All medications have adverse effects. Among the most serious of these are cardiac arrhythmias. Current paradigms for drug safety evaluation are costly, lengthy, conservative, and impede efficient drug development. Here we combine multiscale experiment and simulation, high-performance computing, and machine learning to create a risk estimator to stratify new and existing drugs according to their pro-arrhythmic potential. We capitalize on recent developments in machine learning and integrate information across ten orders of magnitude in space and time to provide a holistic picture of the effects of drugs, either individually or in combination with other drugs. We show, both experimentally and computationally, that drug-induced arrhythmias are dominated by the interplay between two currents with opposing effects: the rapid delayed rectifier potassium current and the L-type calcium current. Using Gaussian process classification, we create a classifier that stratifies drugs into safe and arrhythmic domains for any combinations of these two currents. We demonstrate that our classifier correctly identifies the risk categories of 23 common drugs, exclusively on the basis of their concentrations at 50% current block. Our new risk assessment tool explains under which conditions blocking the L-type calcium current can delay or even entirely suppress arrhythmogenic events. Using machine learning in drug safety evaluation can provide a more accurate and comprehensive mechanistic assessment of the pro-arrhythmic potential of new drugs. Our study paves the way towards establishing science-based criteria to accelerate drug development, design safer drugs, and reduce heart rhythm disorders.

## Introduction

Developing a new drug is an expensive and lengthy process. The estimated average cost to design and approve a new drug is $2.5 billion^1^ and the time to market from the initial discovery into the pharmacy is at least ten years^2^. Many drugs, not just cardiac drugs, interact with specific ion channels in the heart and can induce serious rhythm disorders^3^. Indeed, the major focus of FDA toxicity testing is the pro-arrhythmic potential of a drug, as determined by its effect on repolarization. Specifically, the approval of a new drug requires assessing its impact on the rapid component of the delayed rectifier potassium current in single cell experiments^4^ and on the duration of ventricular activity in animal models and in healthy human volunteers^5^. Unfortunately, the high cost and long time to test new compounds acts as an impediment to the discovery of new drugs^6^. Further, the limited window provided by these criteria onto pro-arrhythmic potential generates false-positives while at the same time preventing many potentially useful drugs from ever reaching the market^7^. Computational modeling and machine learning could significantly accelerate the early stages of drug development, guide the design of safe drugs, and help reduce drug-induced rhythm disorders^8^.

### Torsades de pointes is a serious side effect of many drugs

All pharmacological agents have the potential to impact cardiac repolarization and, with it, the QT interval. The most serious manifestation of both genetic and drug-induced long QT intervals is torsades de pointes, a ventricular arrhythmia characterized by rapid, irregular patterns in the electrocardiogram^9^. Most episodes of torsades de pointes begin spontaneously and revert to normal sinus rhythm within a few seconds; but some persist, degenerate into ventricular fibrillation, and lead to sudden cardiac death, even in patients with structurally normal hearts^10^. In the United States, more than 350,000 sudden cardiac deaths occur each year, but the true incidence of torsades de pointes is largely unknown^11^. Predicting this potentially fatal heart rhythm is challenging given the complex interplay between genetic predisposition and medications, both prescription and over the counter. Increasing evidence suggests that early afterdepolarizations play a critical role in generating of torsades de pointes^12^. Early afterdepolarizations are oscillations during the repolarization phase of the cellular action potential that result from a reduced outward current, an increased inward current, or both^13^. The theory of nonlinear dynamics can help explain the ionic basis of early afterdepolarizations^14^; yet, it remains unclear which ion channels have the strongest effect on creating or suppressing early afterdepolarizations. A better quantitative understanding of the relevant ionic currents would significantly reduce the design space and accelerate drug screening in the early stages of drug development.

### Machine learning could help accelerate drug development

Leading pharmaceutical companies have long recognized the potential of machine learning, especially during the early stages of drug development: On the protein and cellular levels, machine learning can help identify efficient drug targets, confirm hits, optimize leads, and explain the molecular basis of therapeutic activity^15^. On the tissue and organ levels, machine learning can guide pharmacological profiling and predict how a drug that was designed in the lab will affect an entire organ^16^. While using machine learning in the early stages of drug design, target selection, and high throughput screening is almost standard today, the potential of machine learning in the later stages of drug development, toxicity screening, and risk stratification has not been recognized to its full extent^17^. A promising application of machine learning in the context of cardiotoxicity is to combine several experimentally measured and computationally simulated features into a unifying classifier for torsadogenic risk assessment^18^. A recent study demonstrated that a machine learning classifier that combines cellular action potentials and intracellular calcium waveforms provides better torsadogenic risk prediction than one focused on potassium channel block alone^19^. While there is a general agreement between clinical researchers, pharmaceutical companies, and regulatory agencies that computational tools should play a more central role in the pro-arrhythmic risk assessment of new drugs^20^, current efforts focus exclusively on classifiers at the single cell level and ignore ventricular heterogeneity and the interaction of different cell types across the entire heart^21^. We have recently proposed a novel exposure-response simulator that allows us to quickly and reliably visualize how different drugs–either individually or in combination–modulate ion channel dynamics, cellular electrophysiology, and electrocardiogram recordings across ten orders of magnitude in space and time^22^. Combining this simulator with machine learning techniques^23^ would allow us to seamlessly integrate experimental and computational data from the protein, cellular, tissue, and organ scales to assess cardiac toxicity during pharmacological profiling^24^.

Figure 1 illustrates how we use machine learning to combine computational (top) and experimental (bottom) tools and technologies at the single cell (left) and whole heart (right) levels. First, we probe how different ion channels modulate early afterdepolarizations on the single cell level. Using a hybrid computational and experimental approach, we identify the two most relevant channels and systematically screen the two-channel parameter space to quantify the critical blockade that initiates torsades de pointes. Then, we use high performance computing and machine learning to identify the classification boundary between the arrhythmic and non-arrhythmic domains in this space. We validate our approach using computational and experimental electrocardiograms from whole heart simulations and isolated Langendorff perfused hearts. Finally, we demonstrate the potential of our classifier by risk stratifying 23 common drugs and comparing the result against the reported risk categories from the literature.

**Figure 1:**
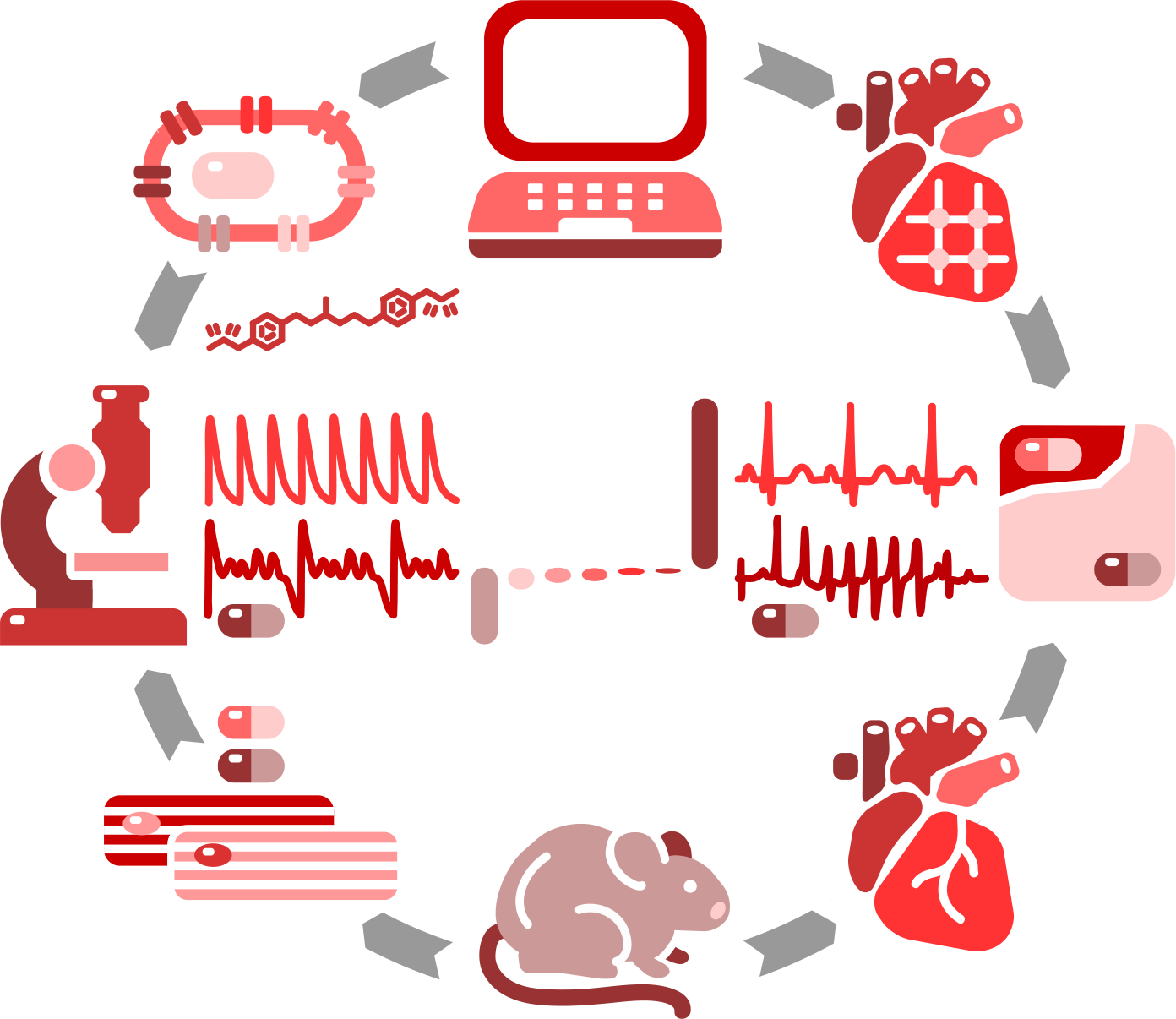
Hybrid computational-experimental approach to quickly and reliably characterize the pro-arrhythmic potential of existing and new drugs. We characterize calcium transients in ventricular cardiomyocytes in response to drugs, both computationally (top) and experimentally (bottom) and identify the ion channels that most likely generate early afterdepolarizations (left). We then screen the concentration space of the two most relevant channels and identify the classification boundary between the arrhythmic and non-arrhythmic domains using high performance computing and machine learning (center). We validate our approach using electrocardiograms, both computationally and experimentally, in whole heart simulations and isolated Langendorff perfused hearts (right). We demonstrate the potential of our new classifier by risk stratifying 23 common drugs and comparing the result against the reported risk categories of these compounds.

## Results

### I_Kr_ and I_CaL_ enhance and prevent early afterdepolarizations

Increasing evidence suggests that early afterdepolarizations are a precursor of torsades de pointes at the cellular level^14^. To identify which ion channels have the most significant impact on the appearance of early afterdepolarizations, we perform 500 simulations of single midwall cells and systematically blocked seven ion channels: the L-type calcium current *I*_CaL_, the inward rectifier potassium current *I*_K1_, the rapid and slow delayed rectifier potassium currents *I*_Kr_ and *I*_Ks_, the fast and late sodium currents *I*_NaP_ and *I*_NaL_, and the transient outward potassium current *I*_to_. Figure 2 illustrates these seven ion channels within the O’Hara Rudy model for ventricular cardiomyocytes^25^. After determining the presence or absence of early afterdepolarizations for all simulations, we fit a logistic regression and extracted the marginal effects, a measure that quantifies the effect of each channel blockade on the probability of early afterdepolarizations. Our results in Figure 2 show that of the seven channels, the rapid delayed rectifier potassium current *I*_Kr_ and the L-type calcium current *I*_CaL_ have the most pronounced effects on early afterdepolarizations. Yet, these two currents display opposite effects: The rapid delayed rectifier potassium current *I*_Kr_ significantly increased the risk of early afterdepolarizations, while the L-type calcium current *I*_CaL_ decreases the risk.

**Figure 2:**
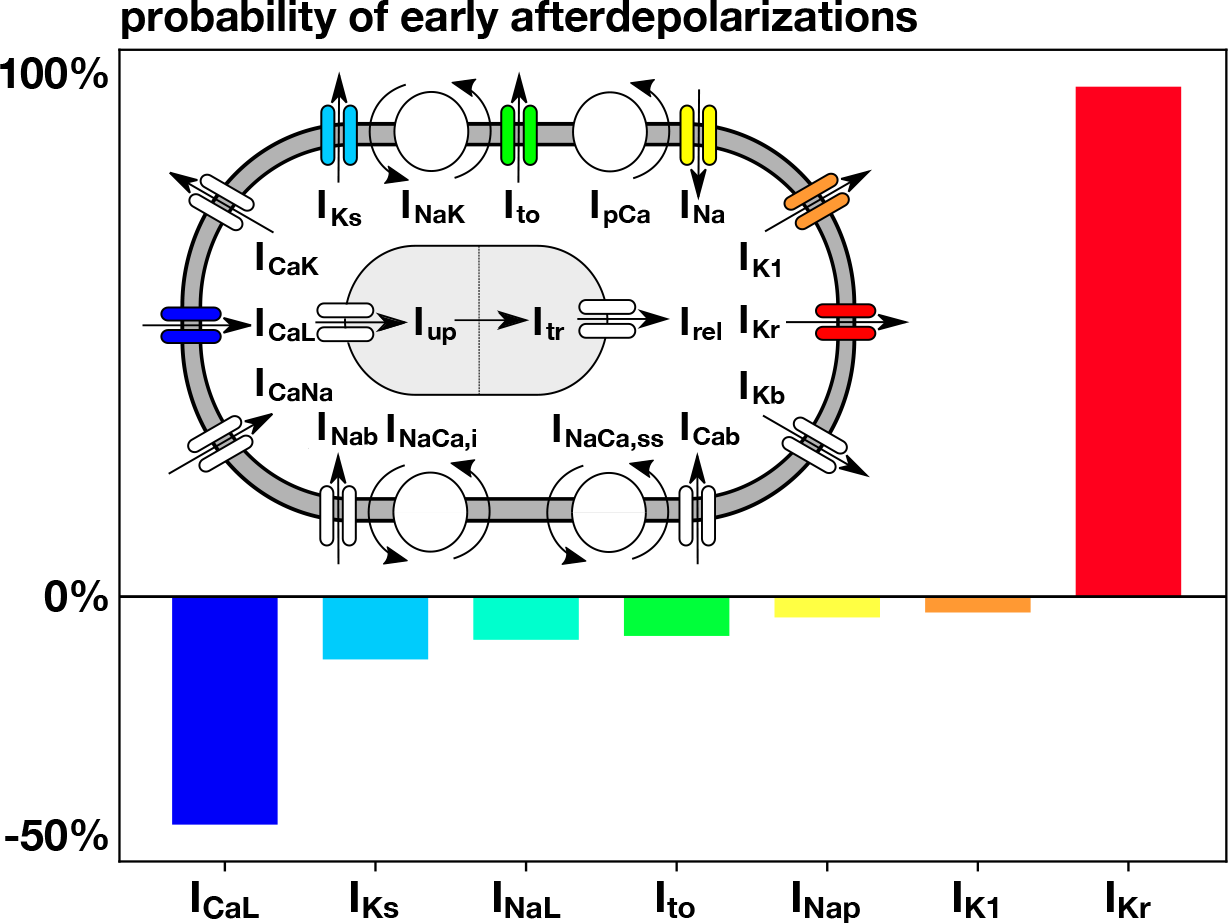
Effect of different ion channels on the probability of early after depolarizations. Positive values imply that blocking this ion channel enhances early afterdepolarizations; negative values imply that blocking prevents early afterdepolarizations. Blocking the rapid delayed rectifier potassium current *I*_Kr_ and the L-type calcium current *I*_CaL_ has the strongest effect on enhancing and preventing early afterdepolarizations.

### I_Kr_ blockade triggers early afterdepolarizations in simulation and experiment

To validate our findings of the computational model, we use isolate rat ventricular cardiomyocytes and expose them to the drug dofetilide, which selectively blocks the rapid delayed rectifier potassium current *I*_Kr_. We record calcium fluorescence and compare it to the calcium transients predicted by the computational model of human ventricular endocardial cells. Figure 3 shows the development of early afterdepolarizations in the presence of the drug dofetilide, both in isolated rat cardiomyocytes and in the single cell model. In both cases, the relationship between the probability of early afterdepolarizations and the concentration of the drug is dose-dependent: Increasing the dose of dofetilide increases the probability of early afterdepolarizations.

**Figure 3:**
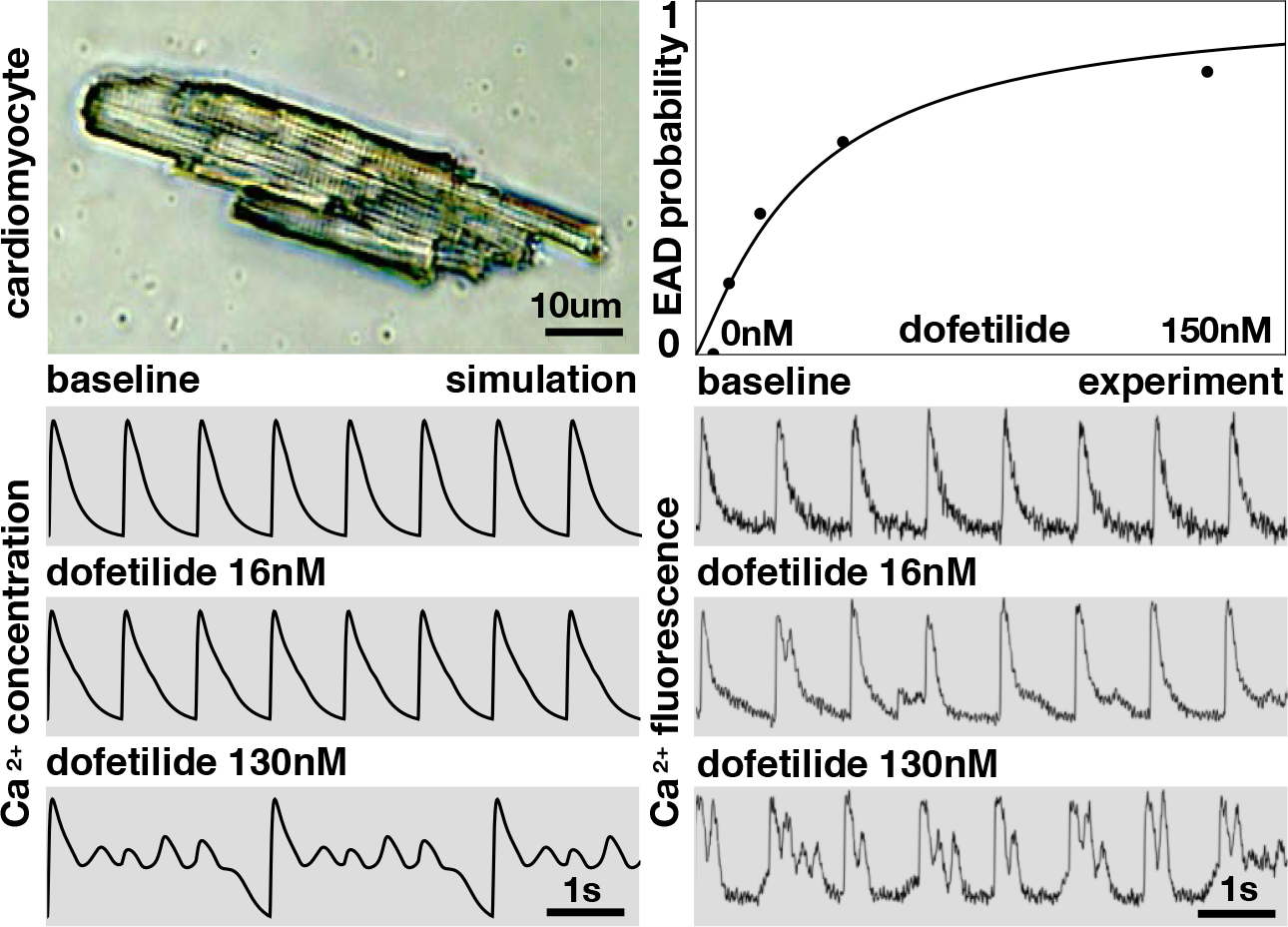
Early after depolarizations in single cell simulation and experiment. Isolated rat cardiomyocyte, top left, and probability to develop early afterdepolarizations in response to the drug dofetilide at concentrations of 4nM, 8nM, 16nM, 38nM, 130nM (n=6 cells each), top right. Calcium transients in response to the drug dofetilide at 0nM, 16nM, and 130nM in the computational simulation, bottom left, and experiment, bottom right.

### Machine learning classifies the boundary beyond which arrhythmias develop

According to our simulated probability of early after depolarizations at the single cell level in Figure 2, we select the two ion channels that most strongly enhance and prevent early afterdepolarizations, the rapid delayed rectifier potassium current *I*_Kr_ and the L-type calcium current *I*_CaL_. We use our high fidelity human heart model^26^ to simulate the effect of combined *I*_Kr_ and *I*_CaL_ block at different concentrations^22^. Our human heart model has 7.5M global degrees of freedom and 0.3G internal variables and runs 1.0M time steps for a simulation window of 5s, which typically takes 40 hours using 160 CPUs. To alleviate the computational cost, we turn to machine learning techniques and adopt a particle learning Gaussian process classifier with adaptive sampling to efficiently explore the parameter space. We randomly perform the first ten simulations and then adaptively sample the points of maximum information entropy determined by our classifier. Figure 4 summarizes the results of our pro-arrhythmic risk classification. The blue electrocardiograms were sampled at points in the blue region and display normal sinus rhythm. The red electrocardiograms were sampled at points in the red region and spontaneously develop torsades de pointes. The white contour indicates the classification boundary. The vertical axis reveals the pro-arrhythmic risk for a selective block of the rapid delayed rectifier potassium current *I*_Kr_: At a critical *I*_Kr_ block of 70%, the risk classification changes from low, shown in blue, to high, shown in red, and the heart will spontaneously develop torsades de pointes. Moving horizontally to the right modulates the pro-arrhythmic risk for a combined block with the L-type calcium current *I*_CaL_: When combining *I*_Kr_ and *I*_CaL_ block, the critical *I*_Kr_ block decreases below 70%. Strikingly, beyond an *I*_CaL_ block of 60%, the heart will not develop fibrillation, no matter how high the *I*_Kr_ block. In agreement with our observations on the cellular level in Figure 2, Figure 4 supports the notion that certain channels can have a positive effect and mitigate torsadogenic risk upon rapid delayed rectifier potassium current block.

**Figure 4:**
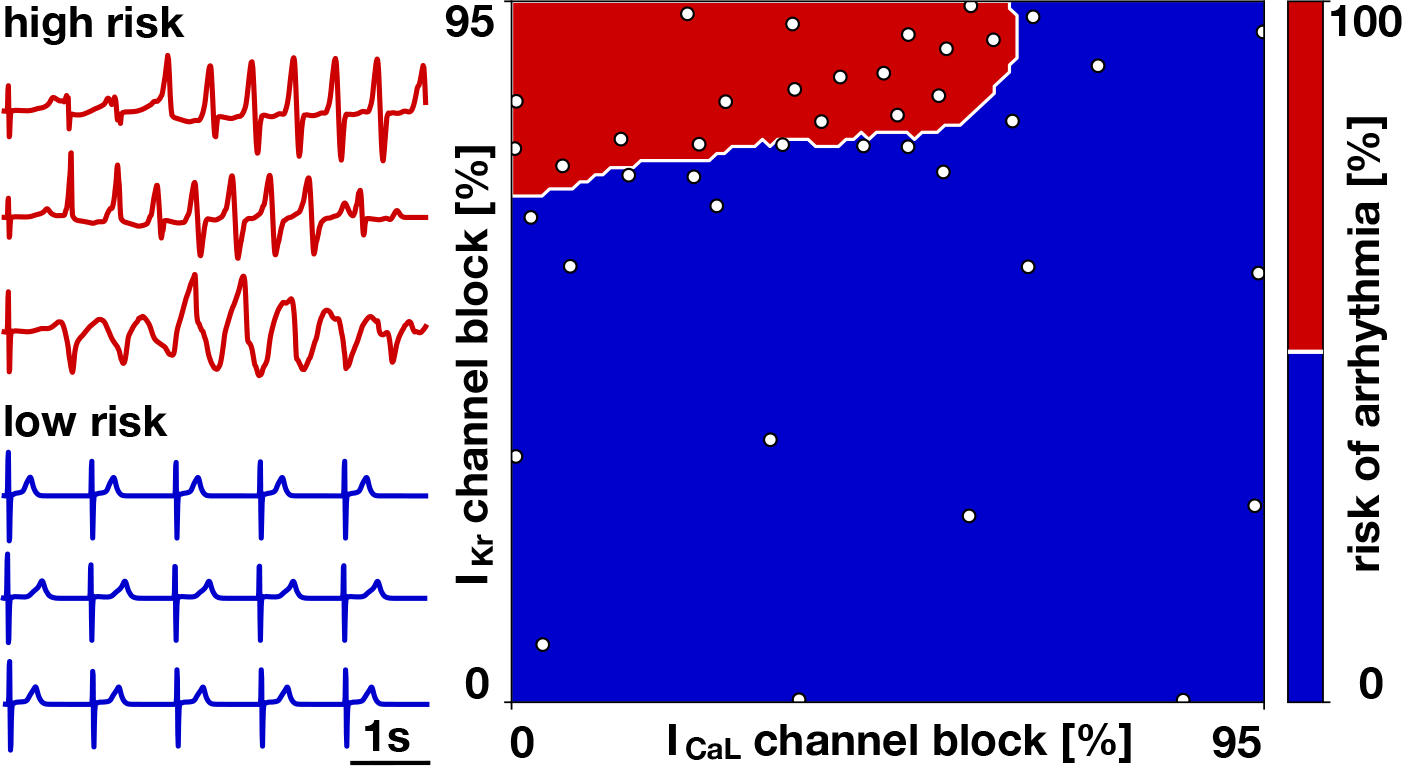
Pro-arrhythmic risk classification. Screening the parameter space of rapid delayed rectifier potassium current *I*_Kr_ and the L-type calcium current *I*_CaL_ block reveals the classification boundary beyond which arrhythmias spontaneously develop. Blue electrocardiograms associated with the blue region displayed normal sinus rhythm; red electrocardiograms associated with the red regions spontaneously developed an episode of torsades de pointes.

### I_Kr_ and I_CaL_ enhance and reduce the risk of ventricular arrhythmias

To explore the interaction between the rapid delayed rectifier potassium current *I*_Kr_ and the L-type calcium current *I*_CaL_ at the organ level, we combine computational modeling and isolated Langendorff perfused rat heart preparations using two different drugs, dofetilide, which selectively blocks the rapid delayed rectifier potassium current *I*_Kr_ and nifedipine, which selectively blocks the L-type calcium current *I*_CaL_. We probe different concentrations of these two drugs and determine the presence of arrhythmias from the computational and experimental electrocardiograms. Figure 5, top, illustrates our Langendorff perfused heart, our four drug concentrations visualized in the pro-arrhythmic risk estimator, and the risk of pre-mature ventricular contractions and arrhythmias for these four cases. Figure 5, bottom, shows the electrocardiograms in response to dofetilide at 0nM and 20nM combined with nifedipine at 0nM, 60nM, and 480nM both for the computational simulation, left, and the experiment, right. For the baseline case without drugs, both the computational model and experimental system display normal sinus rhythm, first row. Blocking the rapid delayed rectifier potassium current *I*_Kr_ by administering dofetilide beyond a critical concentration induces arrhythmias both computationally and experimentally, second row, an observation that agrees well with the single cell simulation and experiment in Figure 3. Additionally blocking the L-type calcium current *I*_CaL_ by co-administering a small concentration of nifedipine markedly alters the excitation pattern both computationally and experimentally, but still triggers irregular beats. Increasing the L-type calcium current *I*_CaL_ block by co-administering a large concentration of nifedipine removes the risk of arrhythmias both computationally and experimentally, the hearts excite at a regular pattern, however at a slightly different rate than for the baseline case without drugs.

**Figure 5:**
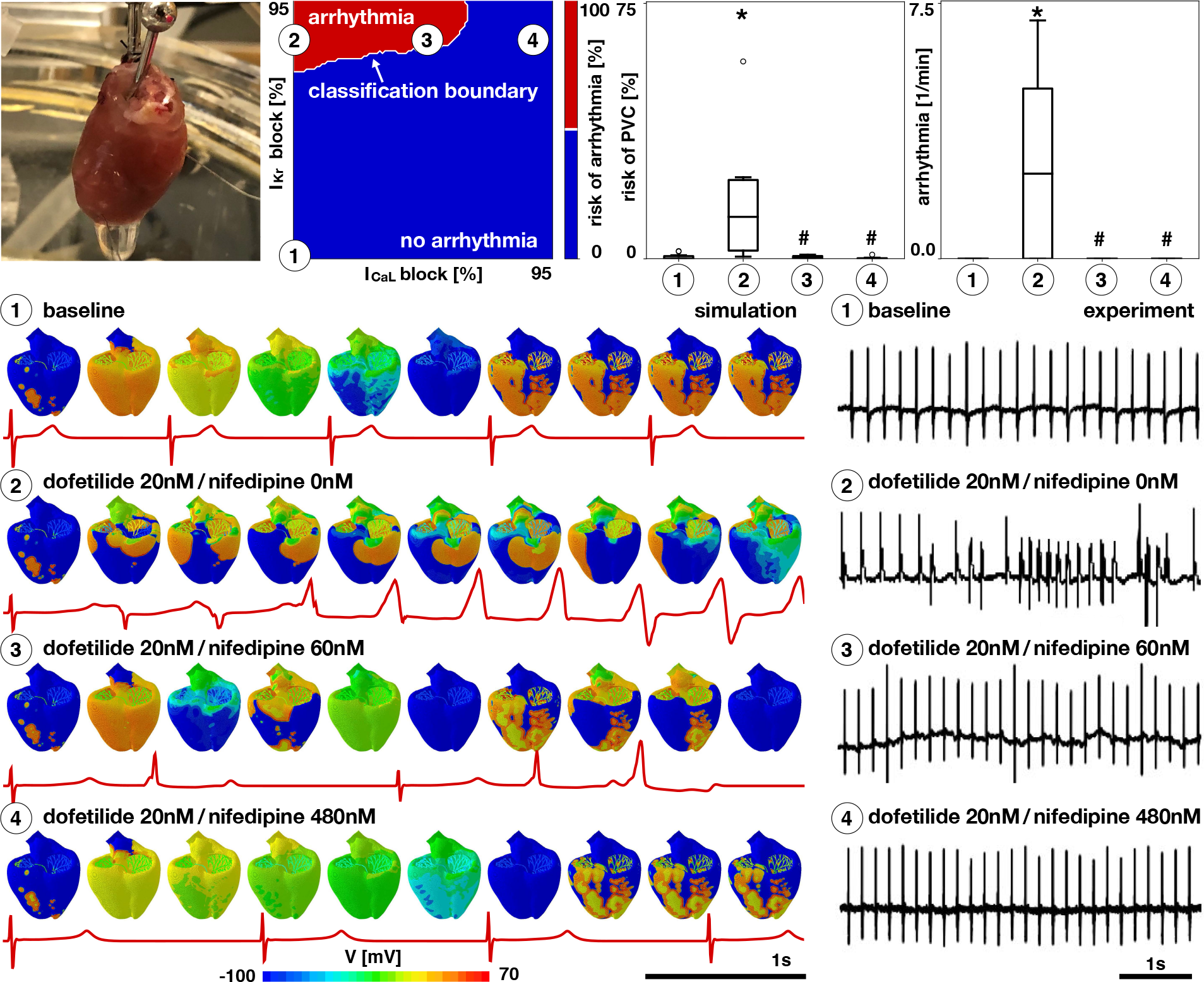
Ventricular arrhythmias in whole heart simulation and Langendorff perfused hearts. Preparation of isolated rat heart, top left, four drug concentrations visualized in the pro-arrhythmic risk classification estimator, top middle, and risk of premature ventricular contractions and arrhythmias in response to varying concentrations of drugs dofetilide and nifedipine (n≥6, * p¡0.05 compared to ➀, # p¡0.05 compared to ➁), top right. Dofetilide selectively blocks the rapid delayed rectifier potassium current *I*_Kr_; nifedipine selectively blocks the L-type calcium current *I*_CaL_. Electrocardiograms in response to dofetilide at 0nM and 20nM combined with nifedipine at 0nM, 60nM, and 480nM in the computational simulation, bottom left, and experiment, bottom right.

### Critical drug concentrations are a predictor of drug toxicity

To validate our approach, we calculate the critical concentrations for 23 common drugs using the risk assessment tool in Figure 4. In essence, the individual block-concentration characteristics for each drug^3, 27^ map onto a trajectory in the *I*_Kr_/*I*_CaL_ plane of the risk assessment diagram. The intersection of this trajectory with the classification boundary defines the critical drug concentration. Curves that never cross the classification boundary indicate a safe drug.

Figure 6 demonstrates that our classification boundary in Figure 4 can reliably stratify the risk of 23 common drugs. Fourteen drugs are classified as high risk drugs. Of those, thioridazine and quinidine cross the classification boundary at the lowest concentrations of 0.1x and 0.3x; chlorpromazine and amiodarone at the highest concentrations of 154.9x and 282.6x. Nine drugs are classified as low risk drugs. Of those, propranolol crosses the classification boundary at 474.6x and all other drugs never cross the classification boundary.

**Figure 6:**
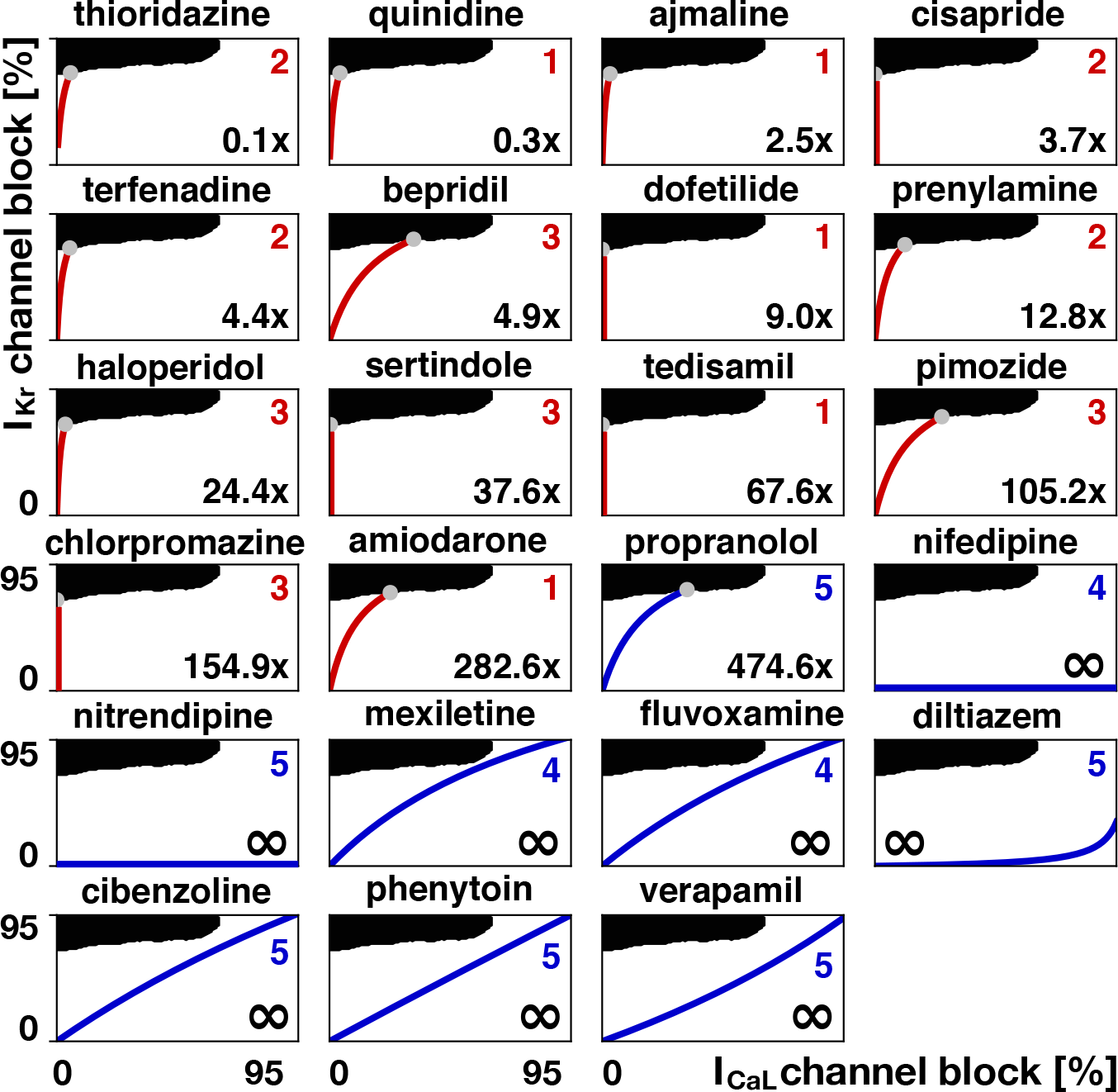
Risk stratification of 23 drugs using our pro-arrythmic risk classification. Black and white regions indicate fibrillating and non-fibrillating regimes; red and blue curves indicate high and low risk drugs; gray dots and numbers indicate the critical concentration at which the curves cross the classification boundary as predicted by our pro-arrythmic risk classification in Figure 4. Numbers from 1 to 5 indicate the reported torsadogenic risk^20^; red and blue colors of the numbers indicate torsadogenic and non-torsadogenic compounds^19^.

Figure 7 illustrates a computational validation of our risk stratification for three drugs, terfenadine, bepidril, and verapamil. Our stratification classifies terfenadine and bepidril as high risk and verapamil as safe. To validate this classification, we apply all three drugs at 10x their effective free therapeutic concentration. Terfenadine, with a critical concentration of 4.4x, triggers an arrhythmia immediately after the first beat; bepidril, with a critical concentration of 4.9x, triggers an arrhythmia after the second beat; and verapamil, which never crosses the classification boundary, is non-arrhythmogenic. While all three drugs initiate a similar degree of blockade of the rapid delayed rectifier potassium current *I*_Kr_ of 84%, 86%, and 79%, their blockade of the L-type calcium current *I*_CaL_ of 11%, 50% and 84% varies significantly. These three examples, now with a complete simulation, highlight the interaction of different channels, and confirm the predictive power of our pro-arrhythmic risk estimator in Figure 4 and its resulting risk stratification in Figure 6.

**Figure 7:**
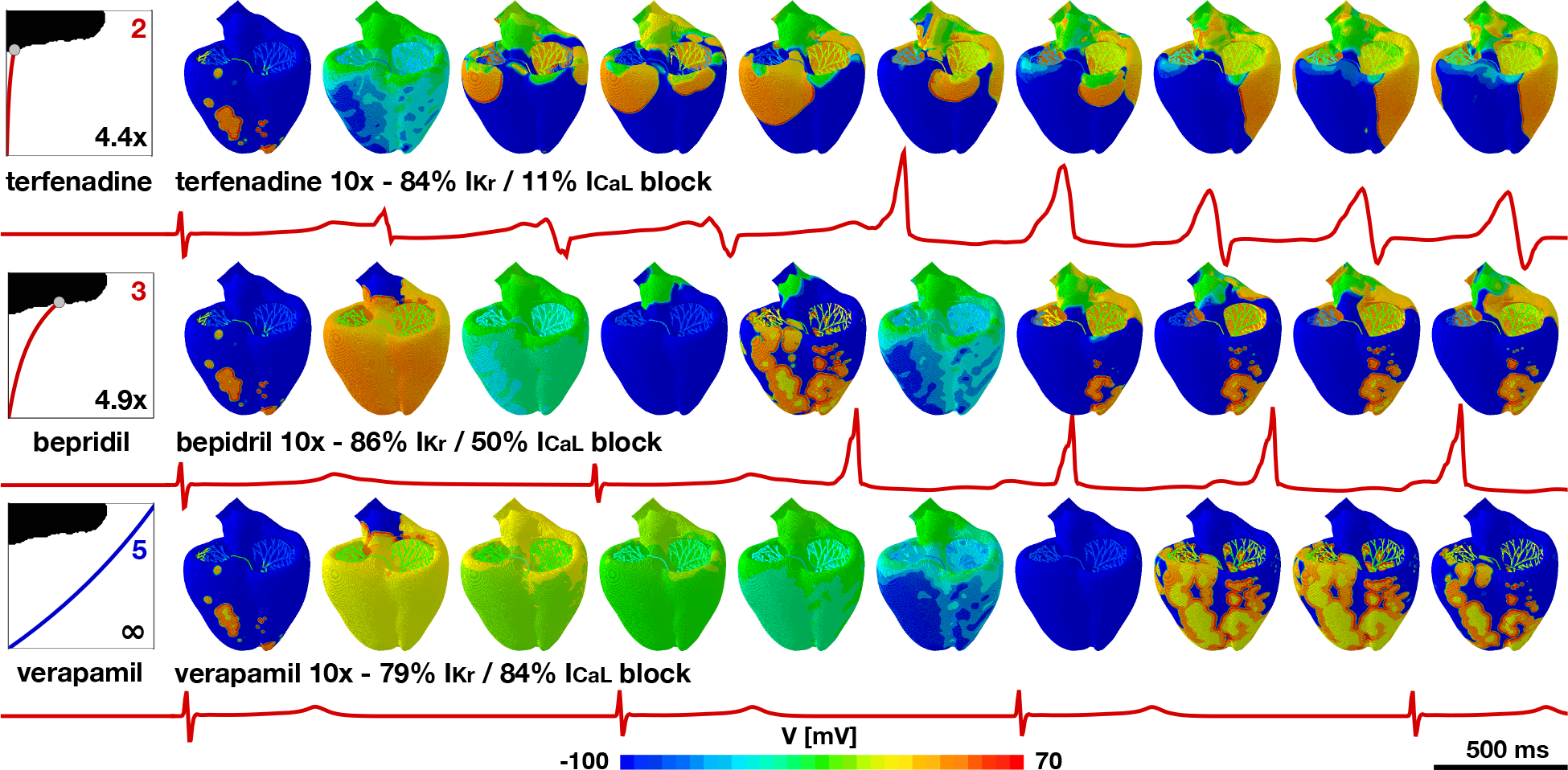
Computational validation of risk stratification for three drugs applied at the same concentration. At 10x the effective free therapeutic concentration, terfenadine blocks 84% of *I*_Kr_ and 11% of *I*_CaL_, bepidril blocks 86% of *I*_Kr_ and 50% of *I*_CaL_, and verapamil blocks 79% of *I*_Kr_ and 84% of *I*_CaL_. The different degrees of blockade result in arrhythmic patterns for terfenadine and bepidril, but not for verapamil, where the high degree of *I*_CaL_ block prevents the development of arrhythmia and slows the beating rate.

## Discussion

Current drug screening paradigms are expensive, time consuming, and conservative. Here we propose a new approach that integrates knowledge from the ion channel, single cell, and whole heart levels via computational modeling and machine learning to reliably predict the cardiac toxicity of new and existing drugs. Our results are based on a rigorous sensitivity analysis that identifies a pair of counteracting ion channels, *I*_Kr_ and *I*_CaL_, that play the most significant role in enhancing and reducing arrhythmogenic risk. We combine multiscale experiments, multiscale simulation, high-performance computing, and machine learning to create a risk estimator that allows us to quickly and reliably identify the pro-arrhythmic potential of existing and new drugs, either in isolation or combined with other drugs. Collectively, these new insights are significant in the development of new compounds. Our efforts significantly extend current initiatives by pharmaceutical industries, clinical researchers, and regulatory agencies with the common goal to develop a new testing paradigm for a more accurate and comprehensive mechanistic assessment of new drugs.

### Early afterdepolarizations are a multiple-channel phenomenon

At the single cell level, we have shown that early afterdepolarizations are triggered when the rapid delayed rectifier potassium current *I*_Kr_ is blocked above a certain level. This is in line with the current regulatory framework, which identifies this channel as the most relevant for QT interval prolongation and torsades de pointes initiation^4^. However, through computational modeling we have demonstrated that early afterdepolarizations are better conceptualized as a multi-channel phenomenon. Our sensitivity analysis in Figure 2 identifies the rapid delayed rectifier potassium current *I*_Kr_ and the L-type calcium current *I*_CaL_ as the most relevant currents for the formation of early afterdepolarizations. These two channels have opposing effects: blocking *I*_Kr_ can initiate and blocking *I*_CaL_ can prevent early afterdepolarizations. In a recent study, we have found a similar trend at the QT interval level^24^, which is also considered in current regulations^5^. These results are in line with other studies that have highlighted the importance of altered calcium dynamics during early after depolarizations^14, 28, 29^, and, more recently, also during delayed after depolarizations^30^. These multi-channel effects between the rapid delayed rectifier potassium current *I*_Kr_ and the L-type calcium current *I*_CaL_ observed in Figure 5 open the door towards a systematic search for blockade combinations that can offset the torsadogenic effects of *I*_Kr_ block alone^31^.

### I_Kr_ and I_CaL_ modulate the onset of torsades de pointes

Our study shows that the rapid delayed rectifier potassium current *I*_Kr_ and the L-type calcium current *I*_CaL_ not only determine the onset of early afterdepolarizations, but also the development of torsades de pointes. Our results in Figure 5 suggest that blocking the L-type calcium current *I*_CaL_ can prevent the development of arrhythmias, even at high levels of rapid delayed rectifier potassium current *I*_Kr_ blockade, both in our high resolution model and in isolated rat hearts. Recent studies have pointed out this preventive role of *I*_CaL_. An analysis of 55 compounds showed that adding the effects of *I*_CaL_ blockade to *I*_Kr_ block improved the predictive potential, while adding the effects of *I*_NaL_ did not^32^. However, this study only demonstrated correlation, without a mechanistic explanation. A recent machinelearning based approach suggested that risk prediction of torsades de pointes could be improved by including intracellular calcium currents^19^. This trend was confirmed by a recent study that classified drugs in terms of *I*_Kr_ and *I*_CaL_ blockade metrics^18^. At the cellular level, these findings reflect the importance of these currents in the development of early afterdepolarizations^14^. At the whole heart level, the presence of these action potential abnormalities is a necessary but not sufficient condition to initiate torsades de pointes; here heterogeneities^33, 34^ and electrotonic effects^12, 21^ play a major role in the propagation of this type of arrhythmia.

### The degree of toxicity correlates with the critical drug concentration

We have classified drugs based on their critical concentration, the concentration at which they cross the classification boundary of our risk estimator in Figure 4. Critical concentration based methods have been used both in rabbit models^35^ and in computational models^36^. Here, we succesfully employed this concept by inducing arrhythmias at elevated drug concentrations both computationally and experimentally. Critical concentrations can be interpreted as the distance from an event of torsades de pointes: The higher the normalized concentration, the further away is the baseline concentration, and thus the safer the compound. When using the critical drug concentration to stratify the risk of drugs in Figure 6, we correctly identify quinidine, bepridil, dofetilide, chlorpromazine, cisapride, and terfenadine as high risk and diltiazem, mexiletine, and verapamil as low risk drugs, similar to a classifier based on net current^37^. Figure 7 confirms the high risk action of terfenadine and bepridil and the low risk action of verapamil, which is widely known as a calcium channel blocker with antifibrillatory effects^38^. Moreover, we correctly identified 22 compounds as high and low risk in Figure 6, compared to the reported high risk categories 1-3 and low risk categories 4-5^20^. For these 22 compounds, our classifier also agrees exactly with a recent machine learning classifier based on action potential duration and diastolic calcium^19^. To eliminate sources of noise in the evaluation of our model, we have only considered those drugs for which 70% or more of the published studies agreed on their risk classification^39^. The only drug that our approach classifies incorrectly is propanolol, which has a critical concentration of 474.6x of the effective free therapeutic concentration. Although Figure 6 suggests that this concentration is significantly higher than for all other high risk drugs, the classifier is trained without any other compound similar to propanolol when performing leave-one-out cross validation. If more data were available, the predictive power of our classifier could be improved. Nonetheless, the potential of our approach lies in supporting the successful progression of compounds that have a poor selectivity to the rapid delayed rectifier potassium current alone and would, under current paradigms, be falsely discontinued through the drug discovery and development process. Our study suggest that our approach correctly identifies those drugs. Our risk estimator in Figure 4 allows us to quickly and reliably screen the pro-arrhythmic potential of any drug, either in isolation or in combination with other drugs.

### Limitations

Although our proposed method holds promise to rapidly assess the risk of a new drug, it has a few limitations: First, our major focus was on combining computational modeling and machine learning to create risk estimators; long term, more experiments will be needed to better validate the method and broaden its scope and use. Second, our model is only as good as its input, the concentration-block curves; we have addressed this limitation in a separate study^24^, similar to other groups^27, 40^, and found that there is a mismatch between the drugs that have been well characterized experimentally^3^–the input of the classifier–and the drugs that we agree in their risk classification–the output of the classifier; to mitigate this limitation, we used a deterministic approach to classify the set of compounds. Third, our current work has mainly followed recommendations of the CiPA initiative^6^; it will be important to validate our model against other cell and heart models, and, probably most importanly, against other compounds. Fourth, we have based our initial studies on reported experiments and clinical observations, supplemented with our own cell level and isolated heart studies with rodent hearts; a critical and logical next step would be to validate our method using our own independent experiments with human adult cardiomyocytes, in larger animals, and, ideally, in healthy human volunteers. Ultimately, with a view towards precision cardiology, our approach has the potential to combine personalized the block-concentration characteristics and personalized cardiac geometries towards identifying the optimal course of care for each individual patient^41, 42^.

## Methods

All studies were approved by the Stanford Administrative Panel on Laboratory Animal Care and conform to the Guide for the Care and Use of Laboratory Animals published by the National Institutes of Health.

### Simulating action potentials in ventricular cardiomyocytes

We modeled the temporal evolution of the transmembrane potential *ϕ* using an ordinary differential equation,

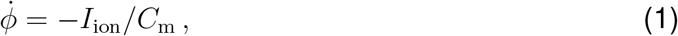

where *C*_m_ is the membrane capacitance and *I*_ion_(*ϕ*, ***q***) is the ionic current, which we represented as a function of the transmembrane potential *ϕ* and a set of state variables ***q***^43^. The state variables obey ordinary differential equations, 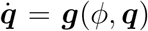, as functions of the transmembrane potential *ϕ* and their current values ***q***^44^. For our single cell simulations, we used ventricular cardiomyocytes with 15 ionic currents and 39 state variables^25^,

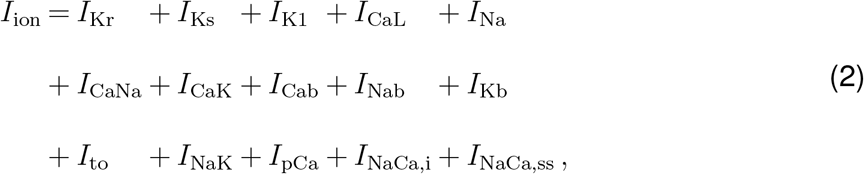

with a minor modification^45^ of the fast sodium current *I*_NaP_^46^. We parameterized the model for human midwall cells^25^, and modeled the effect of drugs by selectively blocking the relevant ionic currents *I*_ion_^47^. For a desired concentration *C*, for each current *i*, we calculate the fractional block *β*_i_ using a Hill-type model parameterized with data from patch clamp electrophysiology^20, 27^, and scale the ionic current *I*_i_ by this fractional block^22^,

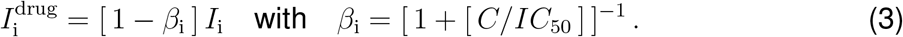

We studied the relative importance of seven ion channels, *I*_CaL_, *I*_K1_, *I*_Kr_, *I*_Ks_, *I*_NaL_, *I*_NaP_, and *I*_to_ on inducing early afterdepolarizations. To achieve a steady state, we paced the cells for 600 cycles at a frequency of 1Hz. We defined the presence of early afterdepolarizations as the occurrence of a change in potential greater than 0.1mV/ms between the 50 and 1000ms of the last two recorded cycles^21^. We used a latin hypercube design to perform 500 simulations and systematically varied the block of the seven ion channels between 0 and 95%. Then, we labeled the results depending on the presence or absence of early afterdepolarizations. We fit a logistic regression and computed the marginal effects, which correspond to the derivative of the output of the regression with respect to the ion channel block. We normalized the results by the maximum value.

### Simulating electrocardiograms in human hearts

To pass information across the scales, we created an ultra high resolution finite element model of the human heart^22^ that represents individual ion channel dynamics through local ordinary differential equations at the integration point level and action potential propagation through global partial differential equations at the node point level^48^. The basis of this model is the classical monodomain model that characterizes the spatio-temporal evolution of the transmembrane potential *ϕ* through the following partial differential equation,

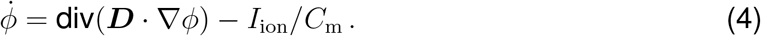

In addition to the local source term *I*_ion_/*C*_m_ from equation (1), the transmembrane potential depends on the global flux term div(***D*** · ∇*ϕ*), where ***D*** is the conductivity tensor that accounts for a fast signal propagation of *D*^‖^ parallel to the fiber direction ***f*** and a slow signal propagation of *D*^⊥^ perpendicular to it^43^,

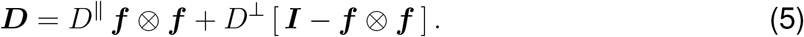

We used the O’Hara Rudy model^25^ from equation (2) for all ventricular cells and the Stewart model^49^ for all Purkinje cells. We discretized the monodomain equation (4) in time using finite differences and in space using finite elements^43^ and introduced the transmembrane potential as a degree of freedom at the node point level and all state variables as local degrees of freedom at the integration point level^44^. We solved the resulting system of equations using the finite element software package Abaqus^50^ with an explicit time integration scheme. We discretized our simulation window of five healthy heart beats in time using 1.0M equidistant time steps of Δ*t* = 0.005ms. We discretized our human heart model^26^ in space using 6.9M regular trilinear hexagonal elements with a constant edge length of *h* = 0.3mm. This results in 7.5M global degrees of freedom and 0.3G local internal variables^48^.

### Using machine learning tools to sample the parameter space

To quickly and efficiently sample the parameter space for a wide range of conditions and a wide variety of drugs we combine our computational models with machine learning techniques^18, 24^. Briefly, to characterize ventricular fibrillation, we performed *n* = 40 human heart simulations and employed a particle learning method to systematically sample the classification boundary within the parameter space. To identify the boundary that divides the arrhythmic and non-arrhythmic domains, we used a Gaussian process classifier and adaptively sampled the point of maximum entropy^51^. We generated the first *n* = 10 samples from a latin hypercube design, and sampled the remaining *n* = 30 samples adaptively. Our results suggest that *n* = 40 simulations are sufficient to reliably identify the classification boundary.

### Classifying drugs into risk categories

We classified 23 drugs into high and low risk, based on our pro-arrhythmic risk estimator in Figure 4 and validated our approach against the known risk classification of these drugs. To select the compounds, we began with a list 31 drugs^20^ for which the concentration block is thoroughly characterized. From these 31 drugs, we only considered those for which 70% or more of the published studies agreed on their risk classification^39,52^, and did not consider the remaining eight controversial drugs. Table 1 summarizes the *IC*_50_ values used to compute the degree of blockade of the L-type calcium current *I*_CaL_ and the rapid delayed rectifier potassium current *I*_Kr_^20^.

**Table 1:** Effect of drugs on ion channels. *IC*_50_ values and effective free therapeutic concentration C_max_ for the 23 drugs used in this study^20^.

| drug | $I_{CaL}$ $IC_{50}$ [nM] | $I_{Kr}$ $IC_{50}$ [nM] | $C_{max}$ [nM] |
| --- | --- | --- | --- |
| ajmaline | 71000 | 1040 | 900 |
| amiodarone | 270 | 30 | 0.3 |
| bepiridil | 211 | 33 | 21.5 |
| chlorpromazine | - | 1470 | 20.5 |
| cibenzoline | 30000 | 22600 | 739 |
| cisapride | - | 6.5 | 3.8 |
| diltiazem | 450 | 17300 | 87.5 |
| dofetilide | 60000 | 5 | 1.2 |
| fluvoxamine | 4900 | 3100 | 196 |
| haloperidol | 1700 | 27 | 2.4 |
| mexiletine | 100000 | 50000 | 2787 |
| nifedipine | 60 | 275000 | 5.4 |
| nitrendipine | 0.3 | 10000 | 1.6 |
| phenytoin | 103000 | 100000 | 4250 |
| pimozide | 162 | 20 | 0.6 |
| prenylamine | 1240 | 65 | 13 |
| propranolol | 18000 | 2828 | 19 |
| quinidine | 15600 | 300 | 2080.5 |
| sertindole | 8900 | 14 | 0.8 |
| tedisamil | - | 2500 | 80 |
| terfenadine | 375 | 8.9 | 4.5 |
| thioridazine | 1300 | 33 | 593.5 |
| verapamil | 100 | 143 | 53 |

### Measuring calcium transients in isolated cardiomyocytes

To characterize calcium transients, we isolated ventricular cardiomyocytes from the hearts of male Sprague Dawley rats with a weight of 250-300g (Charles River, Massachusetts). We anesthetized the rats with inhaled isoflurane and quickly removed the hearts from the chest after euthanasia. We retrograde-perfused the hearts with Ca^2+^-free Tyrode buffer (140mM NaCl, 5.4mM KCl, 0.33mM NaH_2_PO_4_, 0.5mM MgCl_2_, 11mM glucose, and 5mM HEPES at pH7.4) at 1.0ml/min for three minutes, followed by an enzyme solution containing collagenase (1.0mg/ml collagenase type II, Worthington), protease (0.05mg/ml, type XIV, Sigma), and 0.1mM Ca^2+^ for seven minutes. To harvest the cardiomyocytes, we cut the ventricular tissue into small pieces and filtered it with a 250*μ*m nylon mesh. We gradually increased the calcium concentration of the Tyrode solution to 1.0mM for the physiologic analysis and incubated the cardiomyocytes for 15 minutes with 1*μ*M Fura-2-AM (Invitrogen, California) in Tyrode (1.0mM, Ca^2+^). We mounted the cardiomyocytes into a recording chamber on the stage of an Olympus IX-71 inverted microscope (Olympus, New York) where we stimulated them electrically at a frequency of 0.5Hz. Using a galvanometer-driven mirror (HyperSwitch, IonOptix, Massachusetts), we excited Fura-2 at a wavelength of 340/380nm and recorded the emission at 510nm using a photomultiplier (IonOptix, Massachusetts). After five minutes of incubation with the drug dofetilide at concentrations of 4nM, 8nM, 16nM, 38nM, 130nM, we recorded cardiomyocyte calcium fluorescence at 250Hz for eight minutes for n=6 cells each and analyzed the recordings in real time.

### Recording electrocardiograms in perfused Langendorff hearts

To record electrocardiograms, we harvested the hearts of male Sprague Dawley rats with a weight of 250-300g (Charles River, Massachusetts). We excised the hearts from anesthetized rats (2.5% isoflurane in 95% oxygen and 5% carbon dioxide), immediately cannulated the aorta, connected it to a constant pressure perfusion Langendorff system (Harvard Apparatus, Massachusetts) with Krebs solution (118mM NaCl, 4.75mM KCl, 25mM NaHCO_3_, 1.2mM KH_2_PO_4_, 1.2mM MgSO_4_, 1.5mM CaCl_2_, 11mM glucose, and 2mM Pyruvate), warmed to 37° C, and bubbled with 95% oxygen and 5% carbon dioxide. We instrumented the spontaneously beating hearts with ECG electrodes located at the apex and base. After ten minutes of equilibration, we switched the perfusion system to a reservoir to expose the hearts to selected concentrations of dofetilide and nifedipine for a period of five minutes. For n≥6 hearts in each group, we recorded the ECG by Animal Bio Amp (AD Instruments, Colorado) and monitored it continuously throughout the experiment and the a washout period using a Power Lab system (AD Instruments, Colorado).

### Experimentally characterizing the effect of drugs

We characterized the occurrence of arrhythmias in both the isolated cardiomyocytes and the perfused hearts. For the isolated cardiomyocytes, we counted an arrhythmia episode as one if at least one early afterdepolarization occurred within the recording period of eight minutes, and as zero otherwise. We then quantified the relationship between the prevalence of arrhythmia and the concentration of dofetilide using a non-linear regression curve with a two-parameter equation. For the perfused hearts, we calculated the percentage of premature ventricular contractions of all heart beats during the last minute of drug administration. We defined ventricular tachycardia as three or more consecutive premature ventricular contractions. We analyzed the data using Student’s t-test for normally distributed data with equal variance between groups and the Mann-Whitney U test for all other data. For all analyses, we used Prism 7.

## Acknowledgments

This study was supported by the Stanford School of Engineering Fellowship (F.S.C.), the Becas Chile-Fulbright Fellowship (F.S.C.), the Extreme Science and Engineering Discovery Environment XSEDE (E.K.), the Stanford Bio-X IIP Seed Grant program (E.A. and E.K.), and the National Institutes of Health Grant U01-HL119578 (E.K.).

